# Oscillations and aperiodic activity: Evidence for dynamic changes in both during memory encoding

**DOI:** 10.1101/2022.10.04.509632

**Authors:** Michael Preston, Natalie Schaworonkow, Bradley Voytek

## Abstract

Electrical recordings of human brain activity via electroencephalography (EEG) show prominent, rhythmic voltage fluctuations. These periodic oscillations have been linked to nearly every cognitive and perceptual process, as well numerous disease states. Recent methodological and theoretical advances, however, have given rise to evidence for a functional role for non-oscillatory, aperiodic neural activity. Physiologically, this aperiodic activity has been linked to the relative contributions of neuronal excitatory and inhibitory signaling. Most importantly, however, traditional data analysis methods often conflate oscillations and aperiodic activity, masking the potentially separate roles these processes play in perception, cognition, and disease. Here we present a reanalysis of intracranial human EEG recordings from Fellner et al., 2019, using new methods for separately parameterizing oscillations and aperiodic activity in a time-resolved manner. We find that human memory encoding is not related to just oscillations or aperiodic activity, but rather that both processes are rapidly co-modulated during memory encoding. These results provide strong evidence for event-related dynamics of aperiodic and oscillatory activity in human memory, paving the way for future investigations into the unique functional roles of these two independent, but linked, processes in human cognition.

## Introduction

A large body of literature exists that relates neural oscillations to numerous perceptual and cognitive processes, including memory [1,2]. Traditional approaches relate different oscillation frequency bands to distinct components of memory formation, where low-frequency theta (4–8 Hz), alpha (8–12 Hz), and beta (20–35 Hz) have been said to play functionally distinct roles from high-frequency gamma (>40 Hz) [3–7].

In addition to the well-described functional role of oscillations, a new debate has emerged regarding the functional role of non-oscillatory, or aperiodic, neural activity [8–10]. While nascent, there is strong evidence that traditional analysis approaches conflate oscillations and aperiodic activity [10–12]. This likely has far-reaching implications as aperiodic activity has been shown to vary across cortical depth [13], change during development [14–17], and aging [18–20], shift between task states [21], and is altered in disease [22–25]. In particular, the power spectrum slope—or the “spectral tilt”—reflects a global aperiodic neuronal process that is modulated during visual processing [26,27] and working memory [10,28], and is implicated in memory consolidation [29]. The dynamics of this process have been described as a “rotation,” during which broadband power is modulated across all frequencies except a relatively stable rotation point known as the “intersection frequency” that is task-dependent [26]. More recent work, including causal optogenetic manipulation, has physiologically linked the spectral slope and intersection frequency to the balance of neuronal excitation and inhibition [30,31].

Given these new findings, even well-established observations regarding the role of lower frequency (theta, alpha, beta) and higher frequency gamma oscillations in memory encoding have been called into question, where spectral tilts have been proposed as an alternative—or at least complementary—mechanism to multi-oscillation interpretations [7,32] (**Fig 1**).

**Fig 1.**
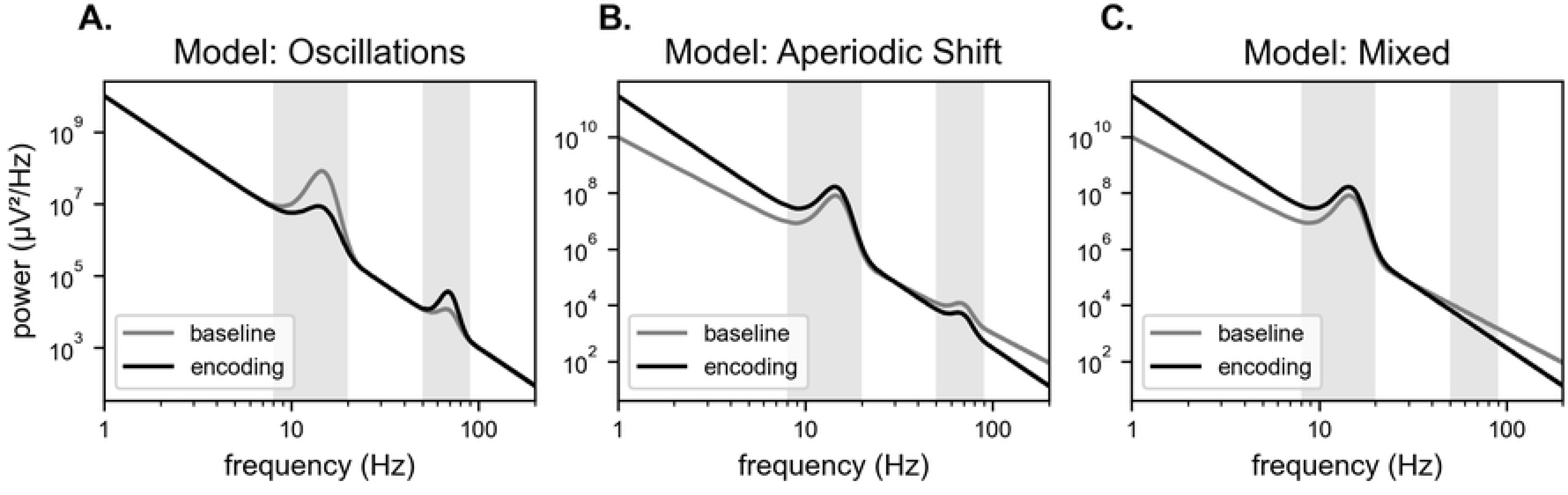
Spectral signatures of memory encoding. Simulated power spectra during baseline (gray) and encoding (black) periods. The low-frequency and high-frequency bands of interest are shaded gray (alpha/beta: 8–20 Hz; and gamma: 50–90). (A) The oscillations (or ‘spectral fingerprints’) model predicts a reduction in the amplitude of a low frequency oscillation and an increase in the amplitude of a high-frequency oscillation. (B) The aperiodic shift (or ‘spectral tilt’) model predicts a decrease in the aperiodic exponent. Importantly, both models predict a decrease in power in the alpha/beta band and an increase in power in the gamma band during encoding. (C) The combined model proposes that encoding is associated with both oscillatory and aperiodic changes. In this example, the spectrum exhibits both a reduction in the amplitude of a low-frequency oscillation and a decrease in the aperiodic exponent.

In a recent series of experiments, Fellner et al. draw a dichotomy between these two models of memory formation, which they refer to as “spectral fingerprints” (oscillations) and “spectral tilt” (aperiodic activity) [33]. The spectral fingerprints model proposes that decreases in lower frequency theta/alpha/beta power and increases in higher frequency gamma power during memory encoding reflect multiple distinct processes (**Fig 1A**); in contrast, the spectral tilt model proposes that these changes in power across multiple frequency bands instead reflect a singular shift in aperiodic activity (**Fig 1B**). Fellner et al. provide evidence that different frequency bands—determined either experimentally (alpha/beta, 8–20 Hz; and gamma, 50–90 Hz) or *a priori* (theta, 2–5 Hz)—exhibit dissociable characteristics with respect to time, space, and stimulus response, arguing against the aperiodic perspective. In a supplemental analysis, Fellner et al. demonstrated that these results were largely stable after correcting for the slope of the power spectrum; however, as the authors note, slope estimates were highly dependent on the frequency band analyzed. Their approach is complicated by the large power deflections caused by the presence of oscillations and nonlinearities in the power spectra such as those from the spectral “knee” [8,10,21,34].

Here, we use methodological advances [10] that explicitly compensate for these features to more fully parameterize the periodic and aperiodic components of the neural power spectra (**Equation 1**). We find that neural power spectra exhibit rapid event-related changes in both oscillatory *and* aperiodic features, simultaneously. That is, rather than a pure oscillatory “spectral fingerprints” model (**Fig 1A**), or a pure aperiodic “tilts” model (**Fig 1B**), there is clear evidence for a mixed model (**Fig 1C**) wherein both processes are dynamically altered during memory encoding. Intracranial electroencephalographic (iEEG) recordings exhibit a dynamic, task-related “tilt”, or rotation about an intersection frequency, caused by a decrease in the aperiodic exponent (i.e., a ‘flattening’ of the power spectrum) following stimulus presentation. This phenomenon occurs in conjunction with a decrease in band-limited, oscillatory power in the alpha/beta band (8–20 Hz). These results show that consideration of both periodic and aperiodic spectral features is essential to our understanding of the neural correlates of memory.

## Results

To quantify oscillatory and aperiodic features, we analyze power spectral content of the baseline (−1–0 seconds) and encoding (0–1 seconds) period by spectral parametrization [10]. This method directly addresses the aforementioned challenges of estimating spectral slope by incorporating spectral peaks and nonlinearities into the model. Specifically, peaks in the power spectrum (i.e., putative oscillations) are modeled as Gaussian functions (**Equation 2**) and removed prior to aperiodic fitting; the aperiodic component is then modeled as a Lorentzian function characterized by a broadband offset, spectral knee, and aperiodic exponent (**Equation 3**). The aperiodic exponent describes the power spectrum slope and is the main focus of the present analysis.

First, we show that both measures, alpha/beta power as well as the aperiodic exponent, decrease during encoding. For each measure, we compared the power spectral content of the baseline (−1–0 seconds) and encoding (0–1 seconds) period at the level of individual electrodes. Data for an example electrode are shown in **Fig 2**; single-trial iEEG time-series can be seen in **Fig 2A** and the power spectra for each time period in **Fig 2B** showing the spectral rotation, characterized by a simultaneous drop in lower frequencies and increase in higher frequencies around an intersection frequency. In the case of the aperiodic exponent, we found 33% and 31% of electrodes exhibit significant decreases (as evaluated by a *p-value* < 0.01 using permutation statistics) within trials of the word and face block, respectively (**Fig 3A**). Furthermore, we find significant decreases in alpha/beta power in 22% and 21% of electrodes within trials of the word and face block, respectively; this includes electrodes exhibiting either an event-related reduction in alpha/beta peak amplitude (as evaluated by a *p-value* < 0.01 using permutation statistics), or a change from measurable alpha/beta in the baseline period to no detectable alpha/beta peak in the encoding period (**Fig 3B**). 9% (word-encoding) and 11% (face-encoding) of electrodes exhibited event-related reductions in both alpha/beta power and the aperiodic exponent. These results demonstrate that oscillatory and aperiodic responses are not mutually exclusive; both measures are modulated during visual memory encoding.

**Fig 2.**
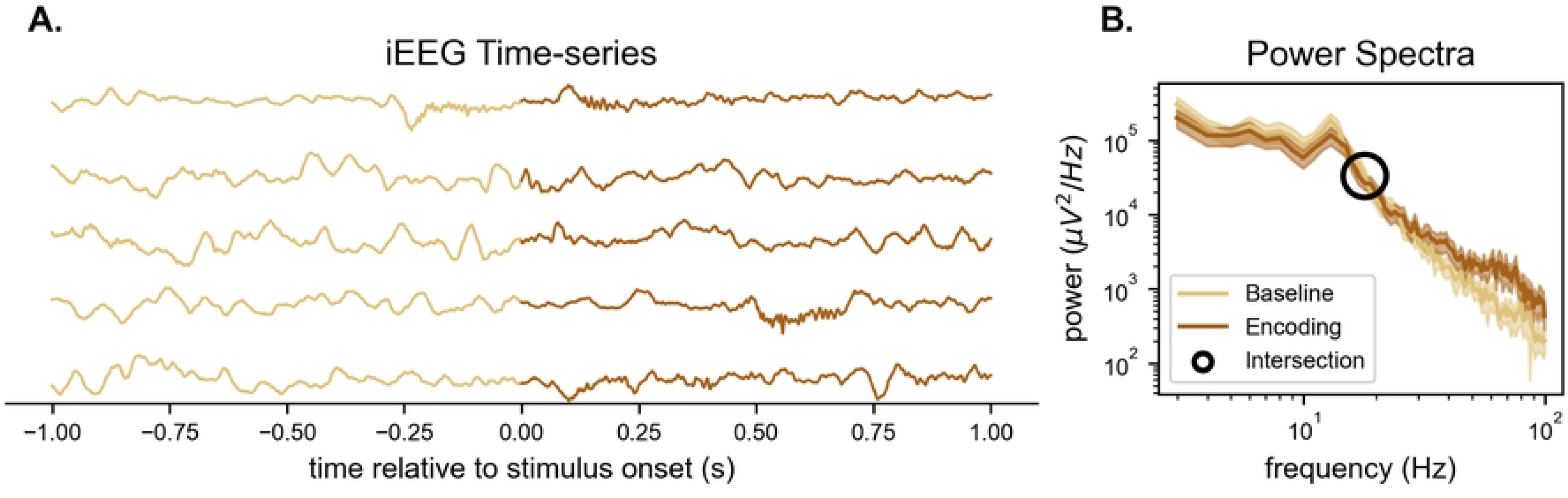
Single-electrode responses to visual stimulation. (A) Single-trial, unprocessed voltage traces for an example recording electrode. (B) Single-electrode power spectra for the baseline (light brown) and encoding (dark brown) period. The across-trial mean is plotted and the 95% confidence interval is shaded. The intersection of the spectra is annotated with a black circle.

**Fig 3.**
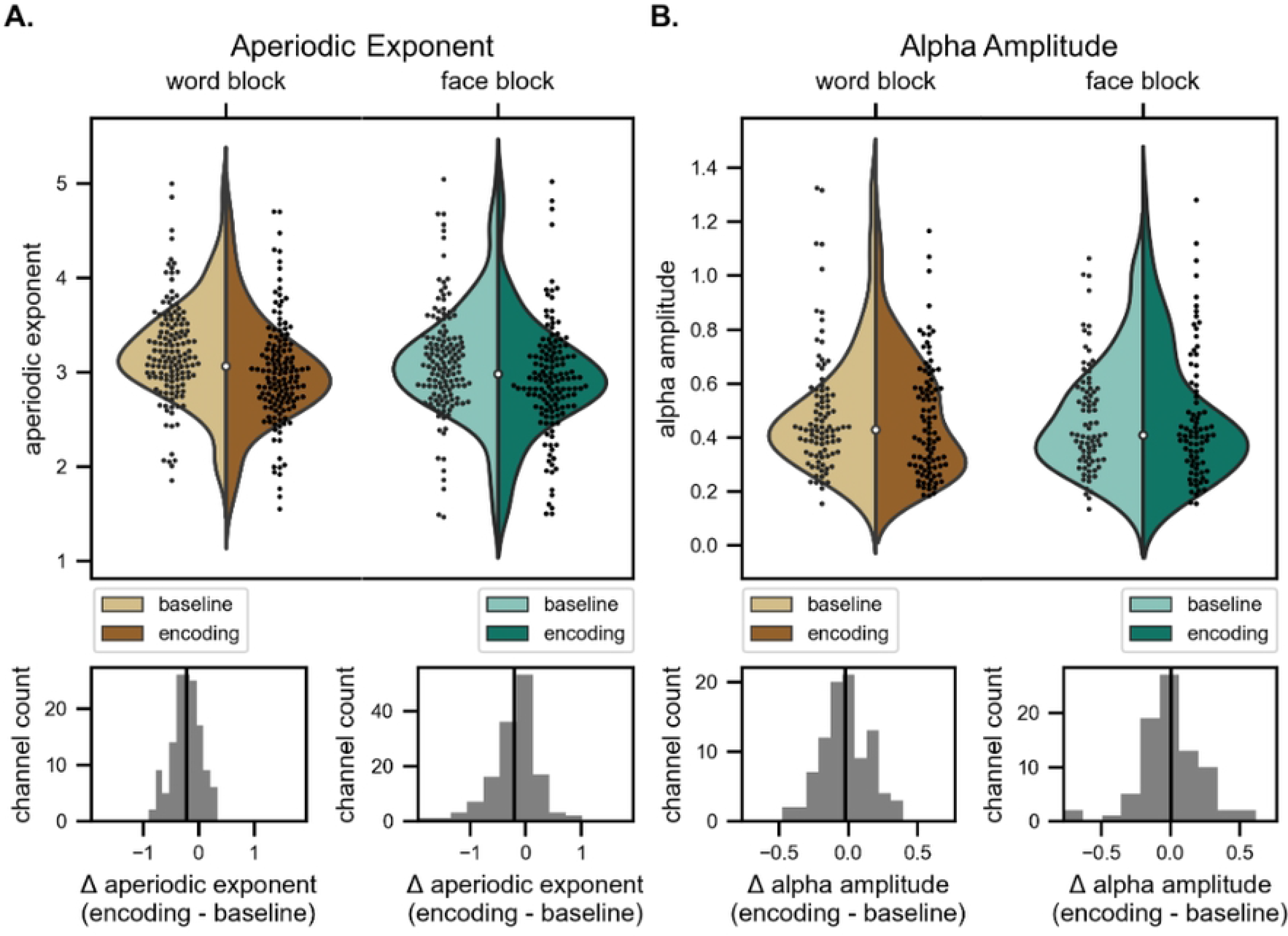
The aperiodic exponent and alpha amplitude are modulated in a task-relevant manner. (A. Upper) Aperiodic exponent during the Word block (left) and the Face block (right). Single-electrode exponents were estimated from trial-averaged power spectra. (A. Lower) Histogram depicting the within-electrode change in exponent between the baseline and ‘encoding period. The black vertical line represents the across-electrode mean. (B) Same as A., except the amplitude of band-limited peaks in the alpha/beta frequency range (8–20 Hz) is plotted. Note that electrodes exhibiting no detectable alpha/beta peak in the baseline period (Word block: 4% and Face block: 7%), the encoding period (Word block: 12%. Face block: 17%), or neither the baseline or encoding period (Word block: 17%. Face block: 17%) are not included in this plot.

To assess the temporal dynamics of these two measures on a more fine-grained scale, we performed time-resolved spectral parametrization, where the measures were extracted for a sliding window over the spectrogram for the whole epoch. An example spectrogram for a single electrode can be seen in **Fig 4A**, as well as the associated measures respectively in **Fig 4B** and **4C**. Averaging across all selected electrodes (**Fig 4D**), it is notable that these changes in alpha/beta power and the aperiodic exponent occur simultaneously, both decreasing rapidly following stimulus presentation with similar temporal dynamics (**Fig 4E and 4F**). This finding suggests that modulation of the aperiodic exponent during encoding contributes to the narrowband results reported by Fellner et al., as total alpha/beta bandpower measures also capture these aperiodic exponent dynamics.

**Fig 4.**
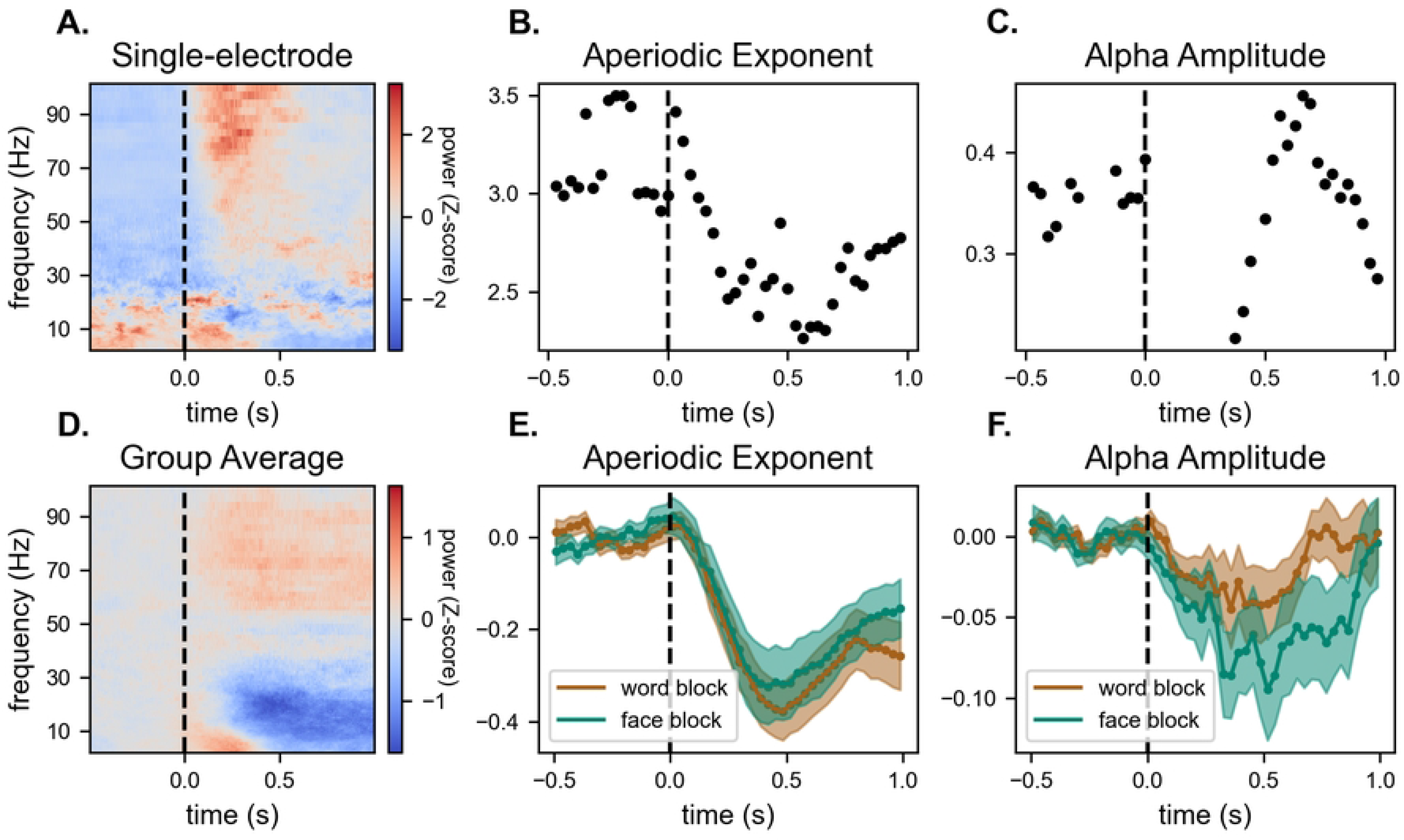
Time-resolved spectral parameterization. Example single-electrode results are shown in A-C, while baseline-subtracted group averages are shown in D-F. (A and D) Spectrograms. Normalized power values represent the across-time z-score, computed individually for each electrode and frequency bin. The vertical dashed line (time=0 s) represents the time of stimulus onset. (B and E) Time-resolved estimates of the aperiodic exponent. (C and F) Time-resolved estimates of alpha/beta amplitude. Note that in some electrodes and at some time points there is no detectable oscillation, resulting in no values given (such as from 0.00 to ~0.25 s in C). Shaded regions in E and F represent 95% confidence intervals.

To investigate how the interplay between aperiodic and oscillatory features can affect conventional bandpower analyses, we next analyzed the intersection frequency i.e., the frequency at which power remains relatively stable during a rotation of the spectrum (**Fig 2B**). This feature is of interest because a change in the aperiodic exponent will have inverse effects on power above and below the intersection frequency. Previous research has demonstrated that power spectra generally rotate around 20–60 Hz during working memory tasks, including in response to visual stimuli (~30 Hz), auditory stimuli (~40 Hz) and motor responses (~60 Hz) (Podvalny, 2015). Consistent with these previous results, we found that, across all electrodes in the full dataset (n=695), the median intersection frequency was 40 Hz, peaking between 10 and 20 Hz; whereas for electrodes with significant effects as reported by Fellner et al. (n=139), the median intersection frequency was 40 Hz, peaking between 20 and 30 Hz (**Fig 5A**). Critically, this frequency is between the high- and low-frequency ranges analyzed by Fellner et al.. This effect was true for both word and face encoding blocks (**Fig 5B**). This finding suggests that decreases in the aperiodic exponent contribute to decreases in low-frequency and increases in high-frequency spectral power, which will change total narrowband power (if not corrected for aperiodic influence), regardless of any changes in oscillatory dynamics.

**Fig 5.**
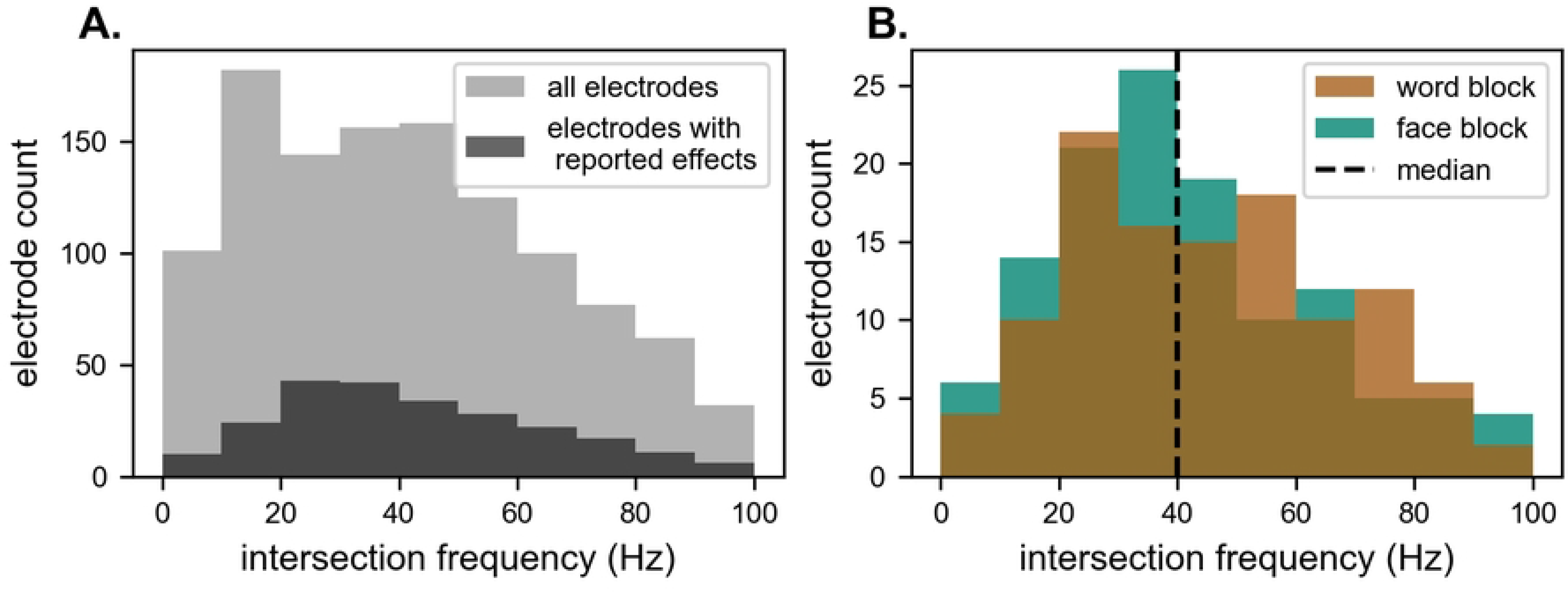
Power spectra rotate about central frequencies. (A) Histogram of the intersection frequency for each electrode i.e., the frequency at which the power spectra for the baseline and encoding period intersect. Electrodes for which Fellner et al. report significant effects are plotted again separately in black. (B) Rotation frequency histograms for the face-encoding block (brown) and word-encoding block (blue) are plotted separately. The vertical dashed line represents the median of all values plotted.

Finally, in order to assess the degree of independence and interdependence between aperiodic activity and oscillations, we performed ordinary least-square regression to evaluate whether shifts in the aperiodic exponent are related to shifts in alpha/beta power, across electrodes. To do this, we regressed alpha/beta power on the aperiodic exponent. We measured alpha/beta power in two ways: in the first approach, consistent with most prior literature, we estimated total bandpower as is traditionally performed, without accounting for aperiodic activity. In this scenario, the hypothesis is that a significant proportion of the event-related change in alpha/beta power can be explained by event-related shifts in aperiodic activity. In the second approach, consistent with the other analyses in this paper that rely on spectral parameterization, we analyzed aperiodic-adjusted peak amplitude in those bands. In this case, we hypothesize that alpha/beta power and aperiodic activity will be more independent.

We find that aperiodic exponent shifts strongly relate to *total* alpha/beta bandpower modulation, estimated using traditional approaches (word trials: *R^2^* = 0.196, *F* = 33.42, *p* < 10^−7^; face trials: *R^2^* = 0.523, *F* = 150.1, *p* < 10^−23^) (**Fig 6A and 6D**). This is consistent with the hypothesis that event-related changes in aperiodic activity can cause what looks like changes in band-limited oscillatory power, even if no oscillation is actually present in the data. In contrast, the aperiodic exponent is not significantly correlated with *aperiodic-adjusted* alpha/beta power (word trials: *R^2^* = 0.041, *F* = 3.866, *p* = 0.052; face trials: *R^2^* = 0.049, *F* = 4.060, *p* = 0.047) (**Fig 6B and 6E**). These findings suggest that, despite the similar time courses for event-related aperiodic activity and aperiodic-adjusted alpha/beta power (**Fig 4E and 4F**), these processes are only weakly correlated on a trial-by-trial basis. In contrast, much of the variance in *total* alpha/beta power can be explained by aperiodic activity alone, but far less of the variance in alpha/beta power is captured by aperiodic activity once that aperiodic activity is accounted for. In other words, while stimulus-evoked modulation of periodic oscillations and aperiodic activity are related, they are mostly independent processes.

**Fig 6.**
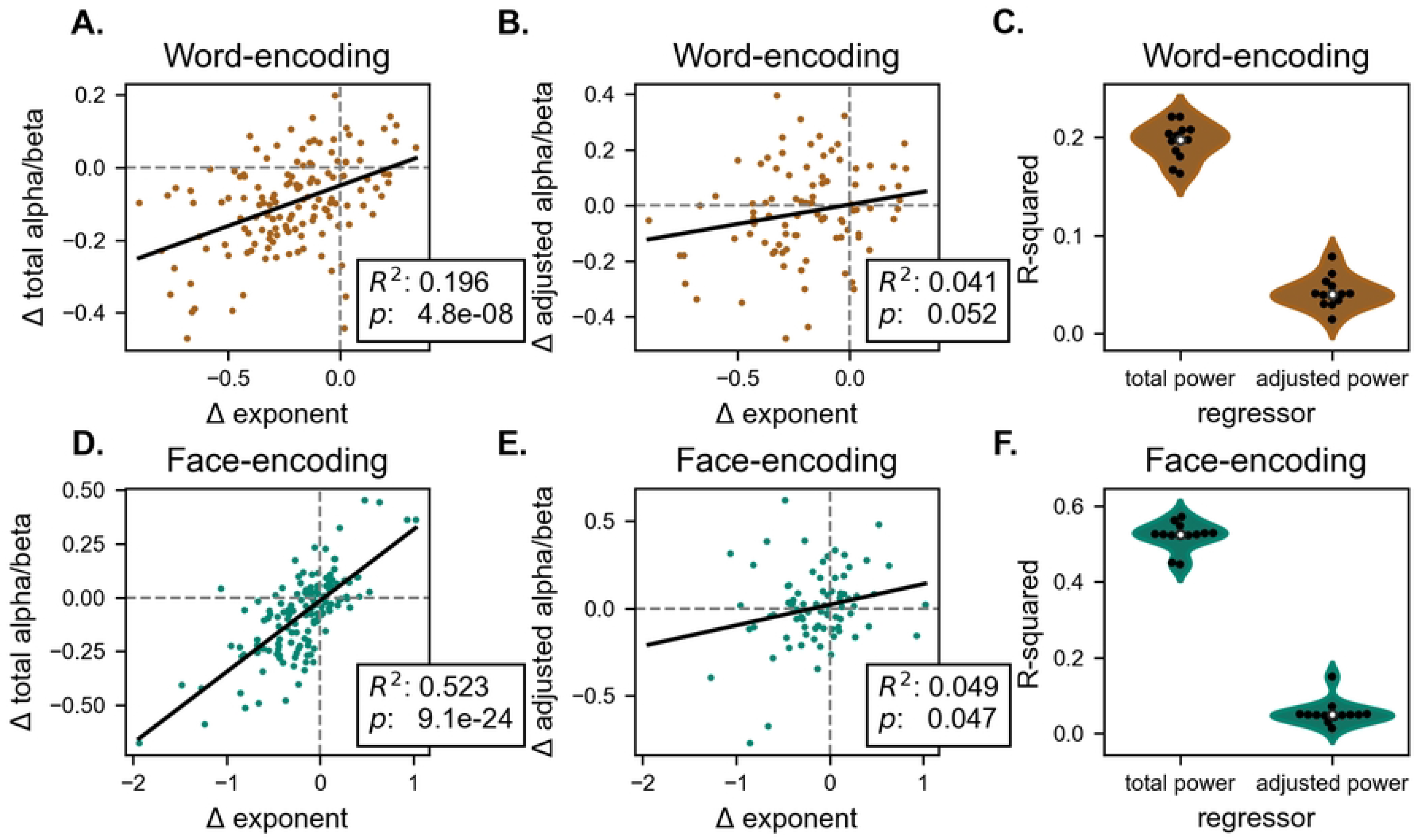
Aperiodic exponent shifts are conflated with total alpha/beta modulation. (A-C) word-encoding block. (A) Scatterplot depicting stimulus-evoked change in *total* alpha/beta power v. the aperiodic exponent. (B) Scatterplot depicting stimulus-evoked change in *aperiodic-adjusted* alpha/beta power v. the aperiodic exponent. (C) Bootstrap analysis results show that the aperiodic exponent explains much more of the variance in total alpha/beta power than does aperiodic-adjusted alpha/beta power. (D-E) Same as A-C, except for face-encoding trials.

Our results show that both aperiodic activity and aperiodic-adjusted alpha/beta oscillation amplitude are systematically modulated during memory encoding. Specifically, the aperiodic exponent decreases, or flattens out, while alpha/beta amplitude tends to decrease.

## Discussion

The present results support the hypothesis that both adjusted alpha power and the aperiodic exponent are modulated during memory encoding. The “fingerprints” *or* “tilt” models are presented by Fellner et al. as dichotomous; however, periodic and aperiodic phenomena are not mutually exclusive. Fellner et al. provide evidence for distinct low-frequency processes during memory encoding, arguing against the aperiodic interpretation [33]. In our reanalysis of their iEEG data, we demonstrate that the aperiodic exponent dynamically decreases—or flattens out—concurrently with the decrease in narrowband, oscillatory alpha/beta power. That is, rather than fingerprints *or* tilts, we show that visual memory encoding is associated with fingerprints *and* tilts.

From a physiological standpoint, the fact that slower oscillations and aperiodic activity are comodulated in human visual cortex may not be surprising due to the importance of inhibitory dynamics in oscillatory formation [35–37] and the suggestive evidence that the aperiodic exponent may partially index excitation/inhibition balance [30,31]. In the context of “high gamma” activity, the event-related “flattening” of the spectrum that we observe here—which has also been observed in human iEEG in the context of visual perception [26]—is particularly striking. While prior work has argued that broadband spectral shifts track population spiking [38–40], the spectral tilt results offer a related, but slightly different, interpretation: instead of population spikes altering total broadband power, task-related shifts in cortical EI balance—driven by excitatory inputs—shift the aperiodic signal to be flatter. This spectral flattening gives the appearance of an increase in high gamma power, along with a simultaneous decrease in lower frequency power, regardless of the presence of oscillations. Each of these phenomena: oscillatory power changes, broadband power shifts, high gamma activity, and spectral tilts, all have different—but likely interrelated—interpretations. With more careful spectral parameterization, we can better link underlying physiology to more complex human cognition.

## Materials and Methods

### Experimental Setup and behavioral paradigm

For this analysis, we reanalyzed data previously collected by Fellner et al. [33]. In the following, we briefly summarize the experimental setup and behavioral paradigm. A more extensive description of the dataset can be found in the original publication [33].

This dataset includes iEEG recordings from 13 patients with pharmaco-resistant epilepsy. iEEG was recorded using a combination of surface grid, strip, and depth electrodes, with a total of 695 electrodes across all patients. Each recording was referenced to a scalp electrode. The sampling rate ranged from 512 to 4,096 Hz.

Neural activity was recorded as patients engaged in a subsequent memory paradigm. Briefly, this task involves an encoding phase in which patients view either words or faces sequentially; and a retrieval phase in which patients are presented with these same items in a pseudo-random order, intermixed with novel items that were not encountered in the encoding phase. The task requires patients to rate their confidence with respect to whether each item was presented during the encoding phase. This classic memory paradigm is commonly leveraged to investigate the neural correlates of memory encoding and retrieval.

### Preprocessing

Fellner at al. applied the following preprocessing protocol to the iEEG dataset leveraged in this study:

1. Segmentation: Data were epoched into trials from −1.5 to 2.5 seconds relative to stimulus onset.
2. Downsampling: Data were downsampled to 512 Hz (to account for the range of sampling rates across patients).
3. Re-referencing: Bipolar montages were computed by re-referencing each electrode to its neighboring electrode (see original publication for more details).
4. Artifact rejection: Data were manually inspected and electrodes with epileptogenic activity were excluded from the dataset.

### Data Analysis

The preprocessed dataset was reanalyzed in Python using open-source toolboxes and custom analysis scripts. Source code to reproduce all analyses shown here from the open dataset is available at: https://github.com/voytekresearch/oscillation_vs_exponent.

The present analysis focuses on electrodes (n=139) for which Fellner et al. report significant differences in bandpower modulation between the word-encoding and the face-encoding block. This contrast was used by the authors to define regions of interest. Furthermore, to specifically isolate spectral signatures of encoding, the present analysis focuses on successful trials only. The average number of trials for the word-encoding block is 62 (SD 19) and for the face-encoding block is 47 (SD 16).

Spectral analyses were performed using the MNE toolbox v.1.0.2 [41]. Power spectra were computed for the baseline (−1–0 seconds) and encoding (0–1 seconds) periods of each trial. The multitaper method was applied to balance the bias-variance tradeoff in spectral estimation of short-time windows [42]. The function mne.time_frequency.psd_multitaper was applied within the frequency range of 2–100 Hz, using a bandwidth of 2 Hz. The average power spectral content for each electrode was then computed as the across-trial mean.

Single-electrode power spectra were parameterized using the spectral parameterization method and toolbox developed by Donoghue et al. v.1.0.1 [10]. In this approach, the power spectrum is modeled as the combination of an aperiodic component and oscillatory peaks:

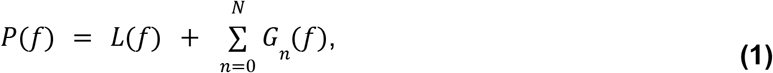

where *P(f)* is the power at frequency *f*, *L(f)* is the aperiodic component, and each *G_n_(f)* is an oscillatory peak. Each oscillatory component, *G_n_(f),* is modeled as a Gaussian function:

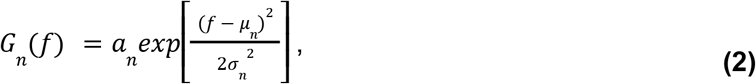

with amplitude *a_n_*, center frequency *μ_n_*, and standard deviation *σ_n_*. The aperiodic component, *L(f)*, is modeled as a Lorentzian function:

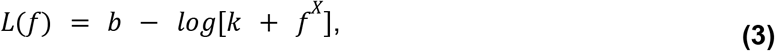

with offset *b*, spectral knee *k*, and aperiodic exponent *X*. This approach improves upon previous methods by explicitly separating oscillatory and aperiodic spectral components and directly measuring the aperiodic exponent or “tilt.” Because aperiodic activity varies between brain regions [21] and across cortical depth [13], aperiodic exponents were estimated individually for each electrode. To reduce the effect of line noise, spectra were interpolated between 45 and 55 Hz prior to parameterization. The following settings were used for spectral parameterization: peak width limits: 2–20 Hz; max number of peaks: 4; peak threshold: 2 standard deviations; aperiodic mode: “knee”. Consistent with the original manuscript, we defined the alpha/beta range to be 8–20 Hz, and restricted our analyses to peaks in this range.

Spectrograms were computed for each trial using multitapers (mne.time_frequency.tfr_multitaper; frequency range: 2–100 Hz; window: 300 ms; bandwidth 10 Hz). The across-trial average spectrogram was then computed for each electrode. Next, single-electrode spectrograms were parameterized using the previously described spectral parameterization toolbox and settings. Prior to parameterization, spectrograms were downsampled to 32 Hz. Model fits were computed for each spectrogram time bin, yielding time-resolved estimates of oscillatory power and aperiodic activity.

The intersection frequency is defined as the frequency at which power remains relatively stable during a spectral tilt/rotation (i.e., a shift in the aperiodic exponent) [26]. The intersection frequency was computed by solving for the intersection of the aperiodic power spectra for the baseline and the encoding period. For each electrode, the aperiodic components of the power spectra were computed (**Equation 3**) using the previously described spectral parameterization results and the NeuroDSP toolbox v.1.65.2 [43].

### Statistics

For each electrode, permutation statistics were used to evaluate the significance of task-evoked shifts in spectral parameters between the baseline (−1–0 seconds) and encoding (0–1 seconds) period. To do this, all spectra from the baseline and encoding periods were pooled, and then were randomly reassigned to be either a baseline or encoding spectrum. For each of these surrogate reassignments, we then compared spectral parameters between pseudo-conditions. This process was repeated (*n* = 1000 shuffling iterations) to estimate a null distribution of task-evoked shifts in each parameter. P-values were then determined based on the probability of each empirical observation given the null distribution derived from the surrogate resampling procedure. An alpha level of 0.01 was used to determine significance.

### Regression

Ordinary least-square regression was used to evaluate whether shifts in the aperiodic exponent were related to shifts in alpha/beta power. For this analysis, both total alpha/beta power and aperiodic-adjusted alpha/beta power were measured. Consistent with the original manuscript, total alpha/beta power was measured as the mean bandpower power between 8 and 20 Hz; consistent with the other analyses in this paper, aperiodic-adjusted alpha/beta power was measured using the spectral parameterization toolbox. An alpha level of 0.01 was used to determine the significance of each model.

Bootstrap statistics were then used to evaluate and compare these regression models. For these analyses, a leave-one-out resampling procedure was applied across patients. For each re-sampled dataset, the regression analysis was repeated and the r-squared value computed to build a theoretical distribution to compare between models.

## Acknowledgements

We thank Fellner and colleagues for sharing their original data (Fellner et al., 2019) and Simon Hanslmayr and colleagues for fun, constructive discussions.

